# Genetic profiling of *Plasmodium ovale wallikeri* relapses with microsatellite markers and whole-genome sequencing

**DOI:** 10.1101/2023.02.01.526392

**Authors:** Valentin Joste, Emma Colard-Itté, Émilie Guillochon, Frédéric Ariey, Romain Coppée, Jérôme Clain, Sandrine Houzé, the French National Reference Center for Imported Malaria Study Group

## Abstract

Like *Plasmodium vivax*, both *Plasmodium ovale curtisi* and *Plasmodium ovale wallikeri* have the ability to cause relapse in humans, defined as recurring asexual parasitaemia originating from liver dormant forms subsequent to a primary infection. Here, we investigated relapse patterns in *P. ovale wallikeri* infections from a cohort of travelers who were exposed to the parasite in Sub-Saharan Africa and then experienced relapses after their return to France. Using a novel set of eight highly polymorphic microsatellite markers, we genotyped 15 *P. ovale wallikeri* relapses. For most relapses, the paired primary and relapse infections were highly genetically related (with 12 being homologous), an observation that was confirmed by whole-genome sequencing for the four relapses we further studied. This is, to our knowledge, the first genetic evidence of relapses in *P. ovale* spp.

## Introduction

Malaria is one of the deadliest infectious diseases, especially in Sub-Saharan Africa with approximatively 619,000 deaths in 2021 [1]. While *Plasmodium falciparum* is the best characterized species in Africa with a well-described epidemiology [1], *P. ovale* spp. are also endemic in Africa [2] but their exact prevalence are still imperfectly established and have been probably largely underestimated [3]. The increased use of qPCR (quantitative Polymerase Chain Reaction) makes the diagnosis of non-*falciparum* or *Plasmodium* coinfections easier [4] and will help in the future to better define the *P. ovale* spp. prevalence in endemic area.

*P. ovale wallikeri* and *P. ovale curtisi* were recently classified as two different species based on distinct genetic patterns [5]. These species are sympatric and display different clinical characteristics that are well-described in imported malaria patients, such as deeper thrombocytopenia and shorter latency period for *P. ovale wallikeri* compared to *P. ovale curtisi* [6,7]. Like *P. vivax*, both *P. ovale curtisi* and *P. ovale wallikeri* have the ability to cause relapse [6,8], defined as recurring asexual parasitaemia originating from liver dormant forms subsequent to a primary infection [9]. Despite *P. ovale* spp. relapses are rarer than in *P. vivax* infections, primaquine has demonstrated its ability to reduce their prevalence in endemic settings [10], but not in non-endemic regions [11]. *P. ovale* spp. relapses are still understudied and the development of genomic tools are necessary to study them. We recently developed a method to selectively amplify *P. ovale* spp. genomes over the human genome [12], permitting whole-genome sequencing (WGS) of these parasites by next-generation sequencing (NGS) from clinical blood samples. This is a promising approach but WGS remains expensive and requires bioinformatics skills not available everywhere. Therefore, a set of easy to type and highly polymorphic markers would complement the genetic tools to study *P. ovale* spp. population genetics.

Microsatellite loci are useful molecular markers to study the population genetics and epidemiology of malaria because they exhibit high level of polymorphism, are generally neutral, and are frequently found in eukaryote genomes. While approximatively 30,000 microsatellites are found in their genomes [13,14], microsatellite typing protocols are missing for both *P. ovale* spp. species, contrarily to other human-infecting *Plasmodium* species [15– 18].

In this study, we investigated relapse patterns in *P. ovale wallikeri* infections from a cohort of travelers who were exposed to the parasite in Sub-Saharan Africa and then experienced relapses after their return to France. Using a novel set of eight highly polymorphic microsatellite markers, we genotyped 15 pairs of isolates from primary (D_0_) and relapsing infections (D_relapse_).

## Methods

### Ethical statement

No specific consent from patients was required since clinical and biological data were collected from the FNMRC database in accordance with the common public health mission of all National Reference Centers in France, in coordination with the Santé Publique France organization for malaria surveillance and care. The study was considered as non-interventional research according to article L1221–1.1 of the public health code in France and only requires the non-opposition of the patient (per article L1211–2 of the public health code).

### Relapse definition & sample collection

A *P. ovale wallikeri* relapse infection (D_relapse_) in the context of imported malaria was defined as asexual parasitaemia recurring 28 days or more after a primary *P. ovale wallikeri* infection (D_0_) treated in France and without any reported trip to a malaria-endemic country between the two infections, thus excluding reinfections. Epidemiological, clinical and biological data were collected through an online patient form completed by French National Malaria Reference Center (FNMRC) correspondents. All isolates’ pairs collected between 2013 and 2021 that fit those requirements were included in the study. We also included *P. ovale wallikeri* isolates from the FNMRC DNA bank to measure the genetic diversity of each microsatellite marker. Genomic DNA was extracted from 200 μL of whole blood using MagNA Pure automaton (Roche diagnostics, United States of America) and eluted in 100 μL. *P. ovale* spp. infections were confirmed using the *Plasmodium* typage kit (Bio-Evolution, France). In-house qPCR-

High Resolution Melting (HRM) was used to differentiate *P. ovale wallikeri* from *P. ovale curtisi* [19].

### Microsatellite development and analysis

We used the PowCR01 reference chromosomes (GenBank assembly accession: GCA_900090025.2) and the software Tandem Repeat Finder [20] to identify genome-wide microsatellites. The criteria for microsatellite selection were restricted to perfect or imperfect microsatellite with unit motifs of 2-10 base pairs (bp) and a minimum threshold of 10 repeats. Once the markers were selected *in silico*, several ones were experimentally tested and further selected based on (i) efficacy of the PCR; (ii) absence of nonspecific bands on agarose gel; (iii) location on different chromosomes; and (iv) easy-to-interpret fragment analysis profiles.

Forward primers were marked in 5’ with 6-FAM or HEX fluorophores (Eurogentec®) and when useful, reverse primers were pig-tailed to reduce non-templated addition of a nucleotide to the 3’ end of the amplicon [21]. We used touchdown-PCR to increase sensibility and specificity of the amplification [22]. The PCR mix, cycle and microsatellite analysis are detailed in the Supplemental methods.

### Measurement of genetic diversity

We measured allele frequencies at each marker using all the isolates (D_relapse_ identical to D_0_ excluded) that were successfully genotyped. We calculated the expected heterozygosity (*H*_*E*_) for each marker using the following formula: 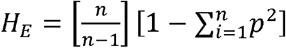 where *p* is the frequency of the *i*^th^ allele and *n* the number of different alleles [23]. We counted the number of alleles (A) at each marker in each country for which more than ten isolates were successfully genotyped. The probability of having two isolates sharing the same allele(s) at all markers was calculated by combining individual probabilities of each marker following this equation: *πp*_*i*_ *= P*_*1*_ *× P*_*2*_ *× (…) × P*_*i*_ where 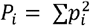 with the assumption of independent infections [17]. The discriminatory power of the microsatellite haplotype was also evaluated with the Simpson index of diversity (*D*): 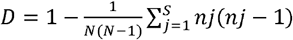 where *N* is the total number of isolates within the population, *s* is the total number of different haplotypes described and *nj* is the number of isolates belonging to the *j*^th^ haplotype.

Polyclonal isolates were defined as isolates with at least two scored peaks for at least one microsatellite marker. Multiplicity of infection (MOI) was calculated for each isolate and averaged across markers.

### Genetic relationship analysis

We built a matrix of the allele count (only the major peak was considered for polyclonal isolates) using the *adegenet* R package [24]. Principal Component Analyses (PCA) were made with *ade4* and plotted with the *factoextra* R package [25]. Phylogenetic tree was built by the neighbor-joining method with the *poppr* R package [26] from a matrix of Bruvo’s distance [27] and plotted with the interactive tree of life (iTOL) software [28].

### Comparison with microsatellite markers of primary (D_0_) and relapse (D_relapse_) infections

We compared the microsatellite haplotypes obtained for the primary and the relapse infections. We classified the two isolates within each pair according to three categories: (i) homologous isolates: all of the alleles detected in the primary infections are detected in the relapse infections; (ii) heterologous isolates: no shared allele(s) between the primary and relapse infections for at least one microsatellite marker; (iii) related isolates: when the primary infection is polyclonal, the D_0_ and the D_relapse_ isolates shared alleles at each marker but some extra-allele(s) is detected only in the primary isolate.

### Selective whole-genome amplification, whole-genome sequencing and variant calling analysis

To confirm the results obtained for relapses with microsatellite typing, genome-wide single nucleotide polymorphisms (SNPs) obtained from WGS were compared for four D_0_-D_relapse_ isolate pairs. Prior to WGS, *P. ovale wallikeri* DNA was enriched following a sWGA strategy and specific primers [12]. Both the sWGA protocol and the WGS procedure are fully detailed in the Supplemental methods.

After WGS, raw reads were mapped to the *P. ovale wallikeri* reference genome (GCA_900090025.2) and duplicate reads were removed using Picard MarkDuplicates (v. 2.26.10). Genome-wide SNPs were identified using BCFtools *mpileup* version 1.13 following previously published quality criteria [12]. We only kept positions with an allelic depth ≥ 5 for the reference and/or the alternative allele. We defined as homozygous a position with ≥ 70% of reads for the reference (homozygous wild-type) or the alternative allele (homozygous mutant) [29]. Otherwise, positions were defined as heterozygous. For genome-wide SNP comparisons, only homozygous SNPs located within the fourteen chromosomes of the nuclear genome were analyzed. Homozygous SNPs located on the contigs and heterozygous SNPs were filtered out.

Genome-wide SNP comparisons were done for each D_0_ and D_relapse_ pair and the proportion of identical genotypes between positions included for both isolates were calculated. We used several WGS data from previous published work [12] or previous unpublished sequencing as controls. The first control was an isolate sequenced twice (either with a sWGA or leukodepleted approach) and named “Matched control pair”. The two other controls were built with one isolate from a relapse pair and one isolate previously sequenced from the same country with no common history and named “Unmatched control pairs”. Finally, to increase the number of comparisons, SNPs from D_0_ and D_relapse_ isolates from different pairs were compared and called “Unmatched infection pairs”.

### Data analysis

All statistical analyses and graphs were performed on R software v.4.0.3 [30]. Quantitative results were expressed as median [Interquartile Interval] or mean +/-Standard Deviation (SD). Spearman’s rank test was used to assess correlations. Proportions were compared using the χ2 or Fisher’s exact tests. We used the Mann-Whitney *U* test or the Wilcoxon signed-rank test to compare medians.

## Results

### Relapse epidemiology

Seventeen pairs of isolates from 2013 to 2021 met the requirements of relapse definition. Patients with relapse mainly lived in metropolitan France and experienced a primary malaria infection after a short travel to Africa (Table S1). Median time between the D_0_ and the D_relapse_ infections was 108 [60-145] days with three relapses that occurred less than 50 days after the primary infection. Delay between D_0_ and D_relapse_ was not different whether the primary infection occurred during high (from July to October) or low transmission malaria season (60 days [46-107] *vs* 121 days [86-142]; *p*=0.19, Mann-Whitney *U*-test) nor in chloroquine or non-chloroquine treated-patients (61 days [50-155] *vs* 128 days [66-142]; *p*=0.39, Mann-Whitney *U*-test). Parasitological follow-up as well as the treatment of the first infections are reported Table S2.

### Sample collection

In order to evaluate the genetic diversity of the selected microsatellite markers, 52 isolates were added in addition to the isolates from relapsing pairs. Those “extra”-isolates were selected to cover several countries in Sub-Saharan Africa and large range of parasite density to evaluate the PCR success rate, especially for low parasite density infections. Sixty-nine isolates were included for microsatellite diversity evaluation (D_relapse_ identical to D_0_ were excluded). They were originated from West or Central Africa (16 different countries) and all were *P. ovale wallikeri* mono-infections (Figure 1). Three isolates were from patients who traveled through multiple African countries (not shown on the map). Ivory Coast (n=15 isolates), Cameroon (n=14 isolates) and Guinea (n=12 isolates) were the most represented countries. Median parasite density was 3,836 [555-9,474] parasites/μL (p/μL).

**Figure 1.**
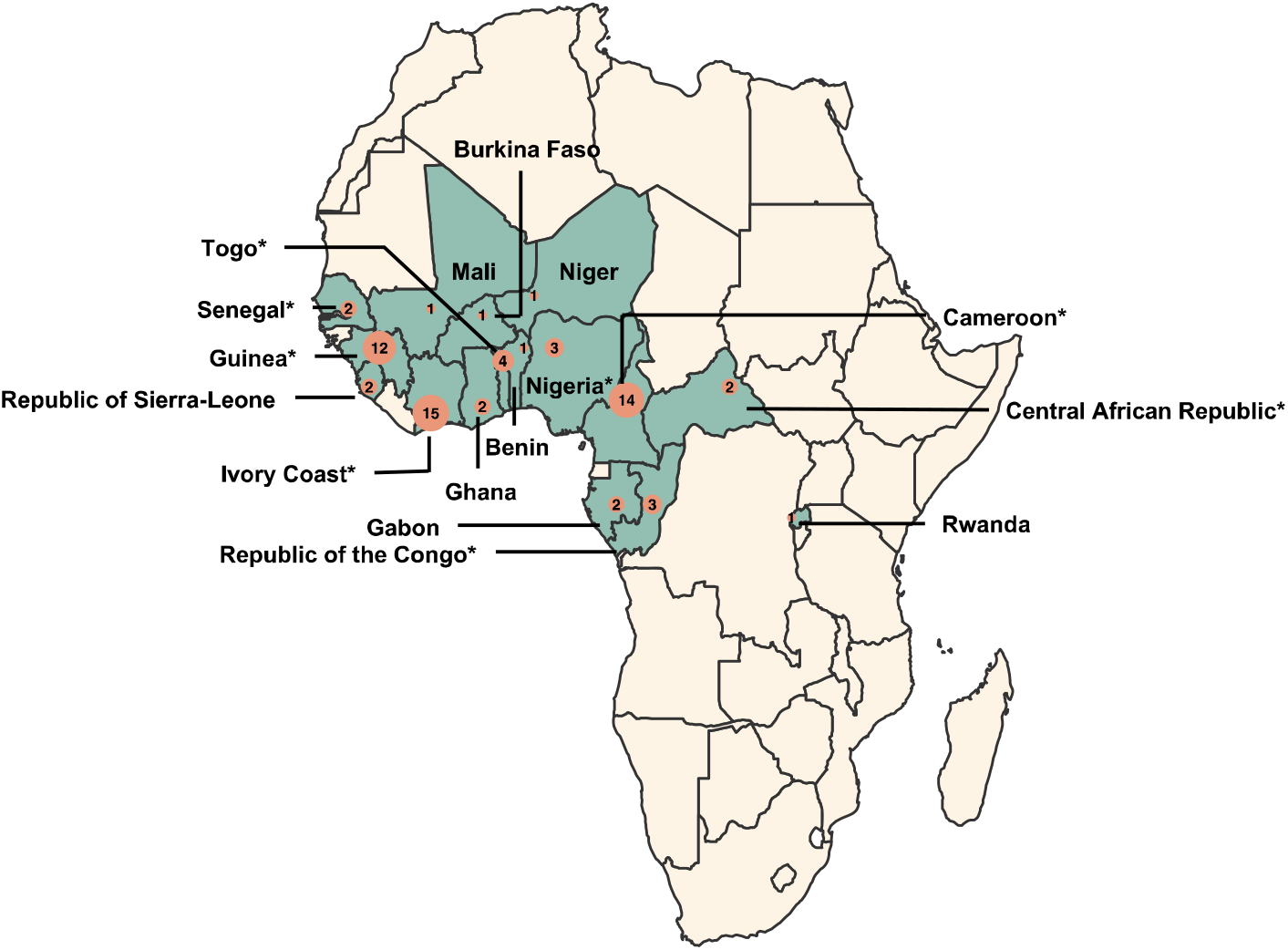
Geographic origins of the *P. ovale wallikeri* isolates included in the study. Countries with included isolates are colored in blue. Number of isolates per country is indicated in salmon circle. * symbols indicate the countries of origin of the relapsing isolates.

### Microsatellite marker development and validation

We developed one-round PCR protocols for eight microsatellite markers, each located on a different chromosome of *P. ovale wallikeri* (Table 1 and Figure S1) and genotyped 69 *P. ovale wallikeri* isolates. Three markers (named 5.2, 11.1 and 14.2) displayed a sensitivity of 100% (no PCR failure; Table 2). The remaining five markers displayed a sensitivity ranging from 94 to 99%. Failed amplifications were associated with a lower parasite density compared to successful amplifications: 6.6% of PCR failures (n=9/136) for parasite densities ≤ 550 p/μL *vs* 1% (n=4/416) for parasite densities > 550 p/μL (*p*=0.001, Fisher’s exact test). Sixty-one samples had a complete haplotype with the eight microsatellites that were successfully genotyped and the remaining had seven (n=5 isolates), six (n=1 isolates) or five (n=2 isolates) microsatellites successfully genotyped. Nonspecific amplification was observed for some markers when tested against *P. falciparum* (marker 2.1) and against *P. malariae* and *P. vivax* (markers 2.1, 8.3 and 14.2). It was significantly decreased when annealing temperature was increased or when DNA templates contained also *P. ovale wallikeri* (Figure S2). As mono-specific *P. ovale wallikeri* infections were analyzed in this study, this caveat had no impact on our results.

**Table 1.**
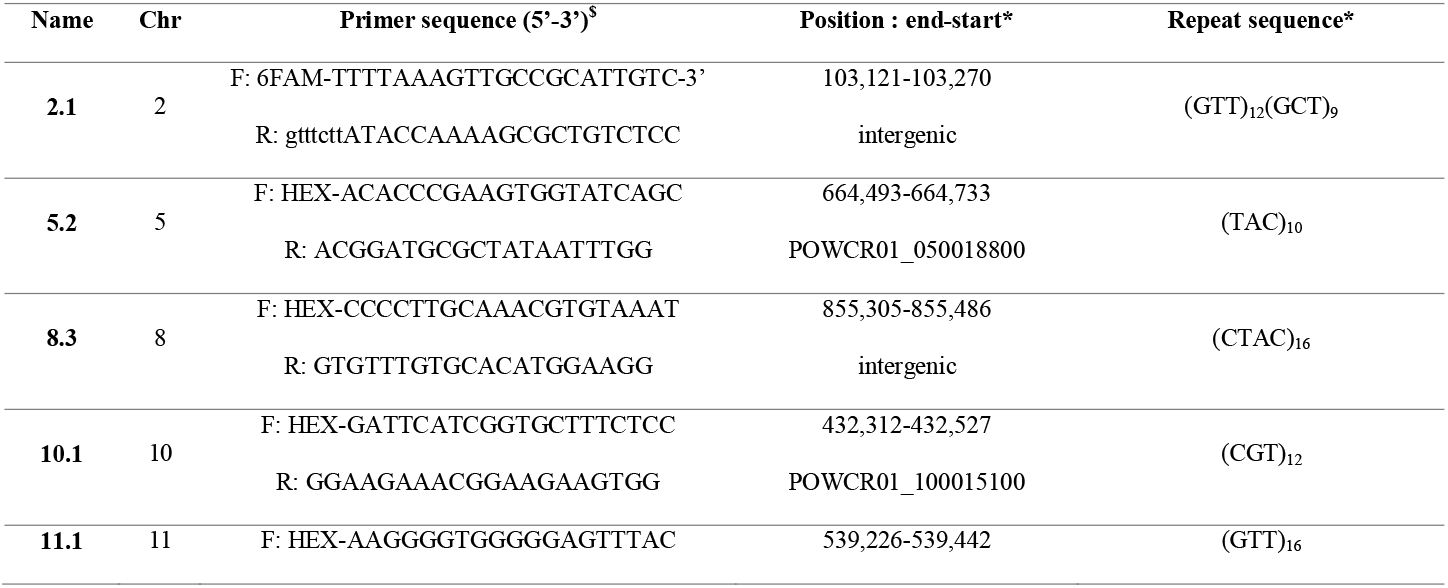
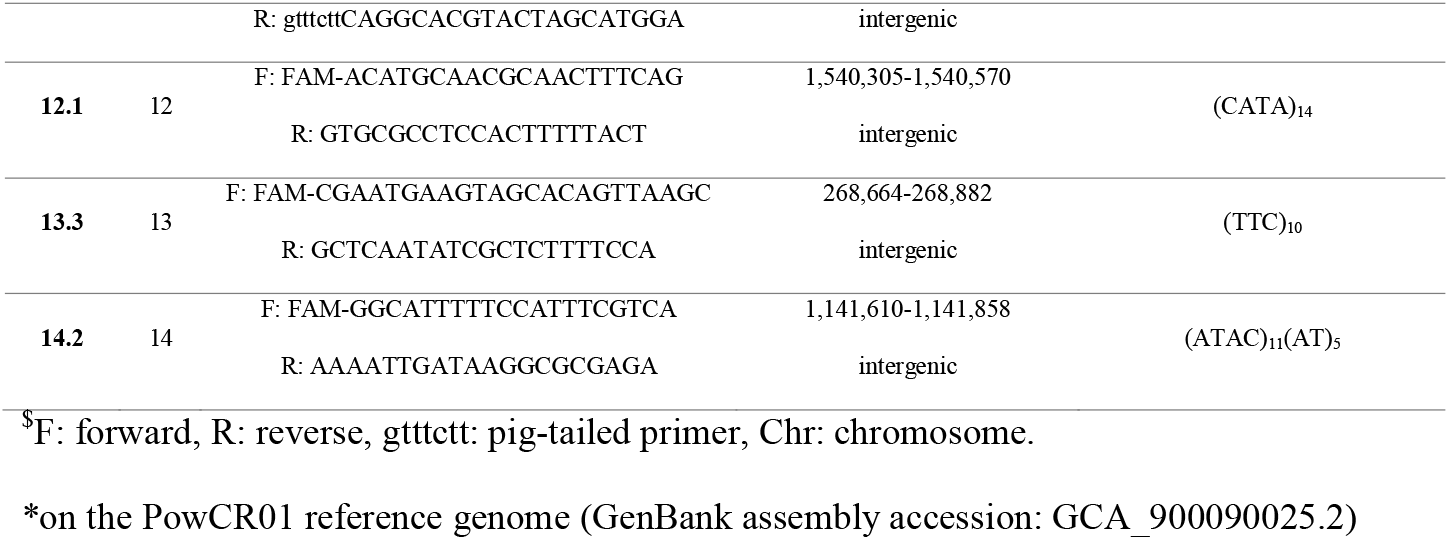
List of the microsatellite markers for *P. ovale wallikeri* genotyping.

**Table 2.** Characteristics of the microsatellite markers developed for *P. ovale wallikeri* (n=69 isolates).

| Microsatellite | PCR length range<br>(bp) | PCR success rate | Allelic diversity,<br>A | Expected<br>heterozygosity<br>(He) | MOI<sup>\$</sup><br>mean (max*) |
| --- | --- | --- | --- | --- | --- |
| **2.1** | 132-159 | 99% | 10 | 0.764 | 1.04 (2) |
| **5.2** | 233-263 | 100% | 9 | 0.849 | 1.01 (2) |
| **8.3** | 127-608 | 99% | 15 | 0.902 | 1.04 (2) |
| **10.1** | 203-239 | 97% | 10 | 0.786 | 1.01 (2) |
| **11.1** | 185-279 | 100% | 17 | 0.932 | 1.03 (2) |
| **12.1** | 227-356 | 94% | 13 | 0.906 | 1.06 (2) |
| **13.3** | 183-267 | 94% | 14 | 0.834 | 1 (1) |
| **14.2** | 227-344 | 100% | 14 | 0.940 | 1.01 (2) |

<sup>\$</sup>MOI: multiplicity of infection.
\*max: maximum of peaks detected in one sample

### Genetic diversity

From the 69 isolates investigated, we counted from 9 to 17 different alleles per microsatellite (Table 2) and 65 eight-marker haplotypes were unique (Table S3). The expected heterozygosity *H*_*E*_ was calculated for each marker and varied from 0.764 to 0.940. For each microsatellite, allele frequencies were calculated in the total cohort (Figure 2 and Table S4) and in Cameroon, Guinea and Ivory Coast (Table S4). The combined likelihood that two random samples had the same eight-markers haplotype was 2.1×10^-6^ (Table S5). The Simpson index of diversity was equivalent (*D*=0.9987) using 8 or only 6 markers (2.1 and 10.1 markers excluded, the two loci with the lower *H*_*E*_). *D* value remained high with the four markers with the highest *H*_*E*_ (*D*=0.9978; combination of the 8.3, 11.1, 12.1 and 14.2 markers), highlighting their combined high discriminatory power.

**Figure 2.**
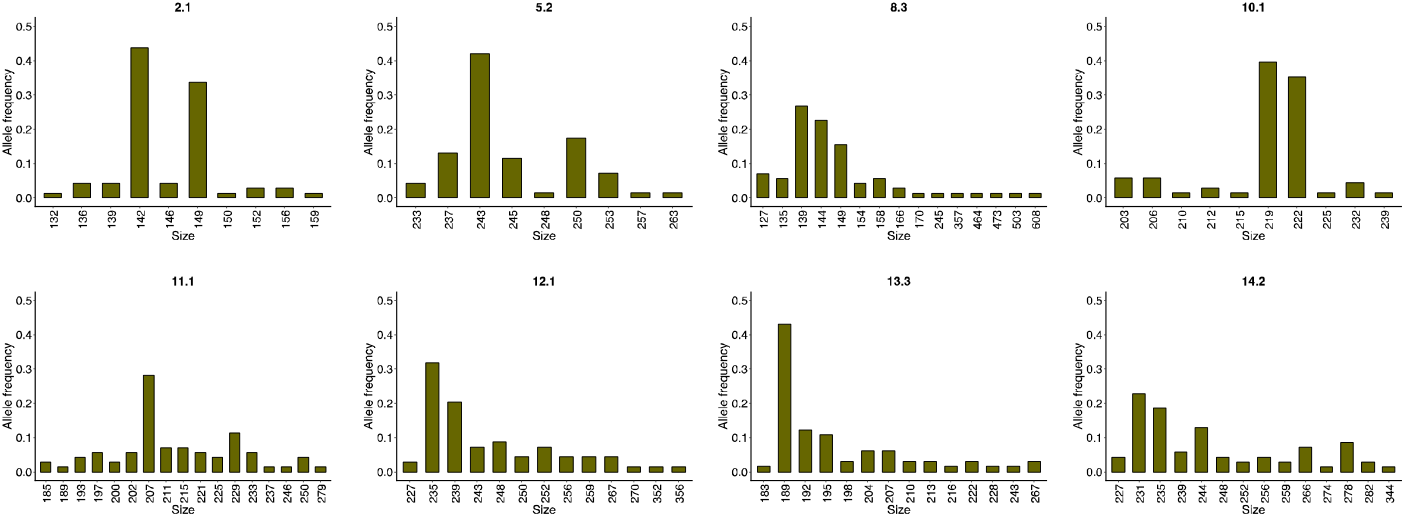
Number of alleles and their frequencies for the eight microsatellite markers.

The diversity (based on the mean number of alleles, A, and on the mean expected heterozygosity, *H*_*E*_) was found to be high in Cameroon and Guinea (Table S6). Interestingly, five of the 69 isolates displayed two peaks for at least one marker and were considered as polyclonal (Table S3).

### Population structure analysis

We next studied population structure between West (n=44) and Central African (n=21) isolates, and between isolates from Cameroon (n=14), Guinea (n=12) and Ivory Coast (n=15), respectively. In both analyses, PCA based on microsatellite allele sizes was not able to separate isolates according to their geographic origin (Figure S3). Similarly, the phylogenetic tree reconstructed using eight-marker haplotypes was not able to separate the isolates from West and Central Africa or from Cameroon, Guinea and Ivory Coast (Figure S4 and S5).

### Application of the microsatellite markers for relapse typing

Among the 17 D_0_-D_relapse_ isolate pairs, two were not analyzable because of PCR failures at more than four markers, likely due to very low parasite density (Table S7). Twelve pairs had homologous parasites (Table S7). Two pairs (pairs 14 and 17) had at least one marker with multiple peaks for the D_0_ but a single shared peak for the D_relapse_ and were then considered as related but not strictly homologous. For one pair (pair 14), the D_relapse_ was monoclonal. For the other pair (pair 17), both D_0_ and D_relapse_ displayed multiple peaks for respectively 4 and 1 microsatellite markers. Finally, the pair 15 had heterologous parasites, with no shared allele for the marker 5.2 (allele sizes were 253 and 250 for D_0_ and D_relapse_ respectively, corresponding to one trinucleotide TAC motif).

### Genome-wide SNP analysis

To complete the microsatellite results, we performed WGS for four homologous D_0_-D_relapse_ isolate pairs (Table S8) and counted the number of SNPs that differed within each pair. For the matched control pair (same isolate sequenced twice), 99.93% of the SNPs were identical, compared to 31.7% for the unmatched control pairs and 30.7% for the unmatched infection pairs (Figure 3 and Table S9). For the four D_0_-D_relapse_ pairs, (pairs 1, 2, 4 and 5; see Table S8) an average of 99.64% SNPs were identical (Figure 3) with very few different SNPs within pairs (Figure 4). No statistical difference in the count of identical SNPs was observed between the matched control pair and the relapses (*p*=0.93, χ2 test). In contrast, the number of identical SNPs statistically differed between the unmatched control pairs or the unmatched infection pairs and the relapses (*p*<0.001, χ2 test).

**Figure 3.**
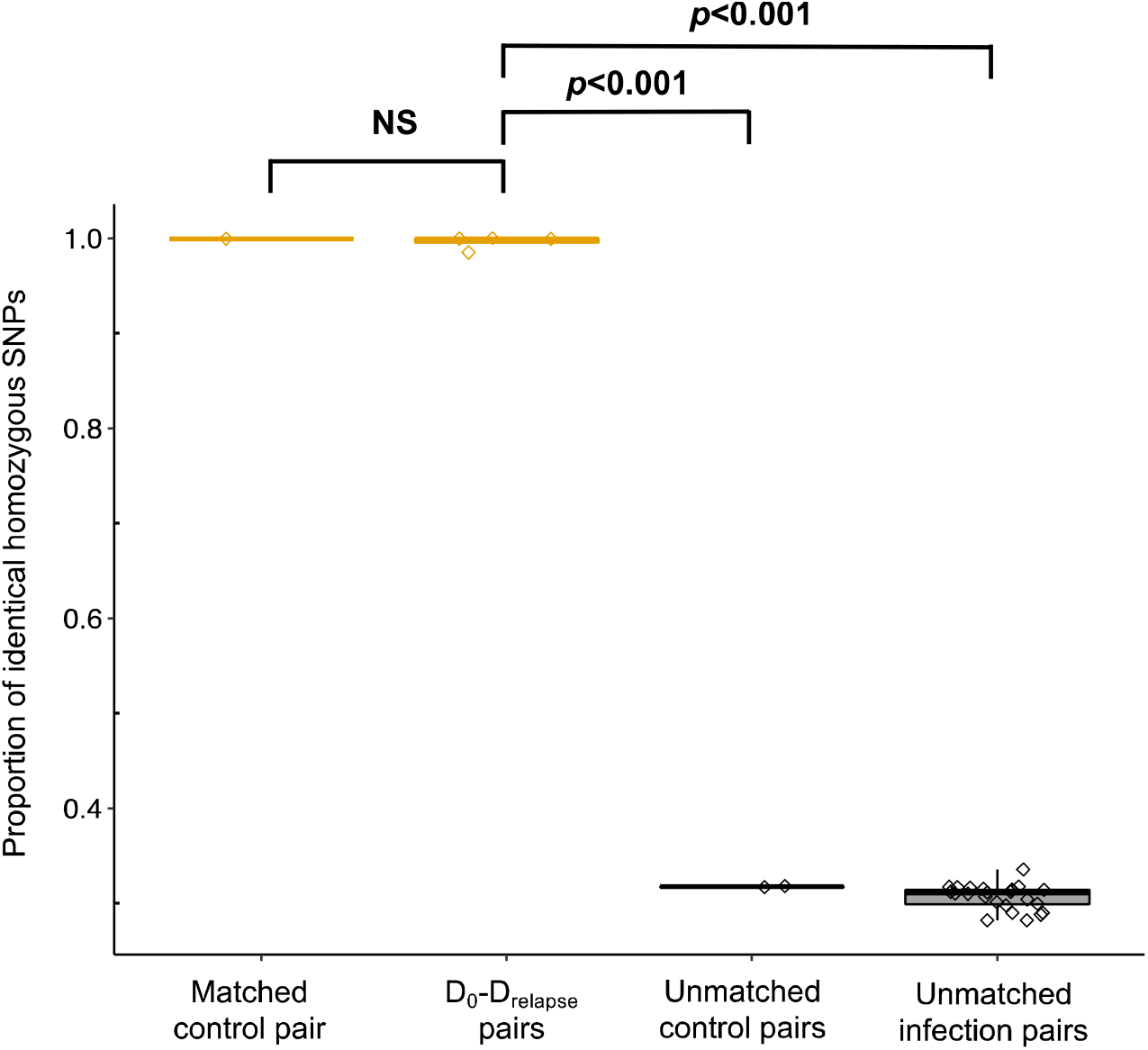
Proportion of identical genome-wide SNPs within infection pairs. Matched control pair and D_0_-D_relapse_ pairs are in light brown. Unmatched control pairs and unmatched infection pairs are in gray. Each dot represents one pair. NS stands for non-significant (χ2 tests).

**Figure 4.**
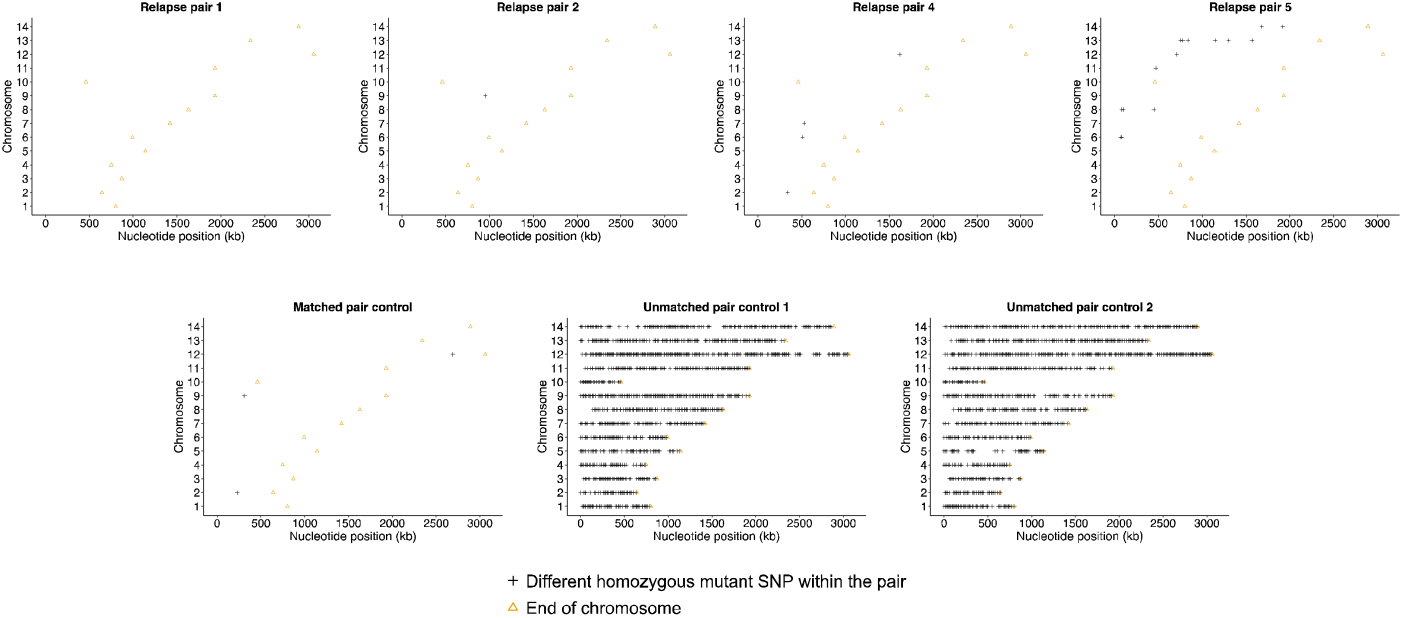
Genetic relatedness within matched or unmatched infection pairs. The distribution of the different alleles within each D0-Drelapse pair is shown throughout the fourteen chromosomes of the nuclear genome. Each cross represents a 10 kb window containing at least one differing SNP, represented according to its genomic position (x axis, nucleotide position in kilobase pair; y axis, chromosome). Beige triangle shows the end of each chromosome.

## Discussion

Despite several epidemiological evidences of *P. ovale* spp. relapses during the last seventy years [6,8,11,31–33], biological realness of *P. ovale* spp. hypnozoites is still debated [9,34]. Indeed, proofs of hypnozoite existence are scarce, but Soulard *et al*. [35] described late-developing schizontes in humanized mice infected with *P. ovale* spp. sporozoites, consistent with a hypnozoite origin of those maturing forms [35].

Easy-to-use molecular tools are missing to study the genetic relatedness of *P. ovale* spp. isolates. We developed eight microsatellite markers that displayed very high *H*_*E*_ and high combined probability value (Table 2), suggesting a great discriminative potential for epidemiological studies. Interestingly, the combination of just four markers (8.3, 11.1, 12.1 and 14.2) remains highly informative. These could be used as a minimal set to genetically profile *P. ovale wallikeri* isolates for time and resource saving. We observed high diversity with 65 unique haplotypes and only 3 non-unique haplotypes each shared by two isolates from West Africa that did not have reported common history (Figure S4 and Table S3). More studies, in particular at a country level, are needed to better investigate *P. ovale wallikeri* population structure in West and Central Africa.

With these eight newly developed microsatellite markers, we successfully genotyped 15 pairs of *P. ovale wallikeri* isolates that fitted with an epidemiological relapse definition. As our study is the first one that genetically analyzed *P. ovale wallikeri* relapses, we used *P. vivax* relapses as a reference. Our relapse definition is purely based on the epidemiology and history of cases as we did not have complete parasitemia follow-up for each patient neither formal evidence of perfect compliance to full treatment. As the World Health Organization recommends a 28 days parasitaemia follow-up for *P. vivax* [36], we hypothesized that infections with time to recurrence above this 28-days duration threshold were relapses. Of note, even with precise molecular tools, a homologous relapse infections cannot be formally distinguished from a recrudescent infection caused by incomplete treatment clearance of the blood-stage primary infection. We decided to classify those second infections as homologous relapse and not recrudescence because: (i) in *P. vivax*, recrudescences seem to be much less frequent than relapses [37,38]; (ii) to the best we know, *P. ovale* spp. treatment failure is rarely reported in the literature (no failure reported in published clinical trials [39–43], except one persistent *P. ovale wallikeri* on day 28 in a *P. falciparum* co-infection in Gabon [44]); (iii) except for three pairs for which the time to recurrence was short (28, 33 and 39 days), this time was much longer for the other pairs (from 53 to 257 days) which is not in favor of a recrudescence. In addition, those patients were not living in endemic areas and therefore had likely not acquired sufficient immunity to maintain for a long time a low-level malaria infection with *P. ovale* spp. A prospective study, with WHO parasitological follow-up, directly observed treatment and plasma drug dosage is however needed to further confirm our conclusions.

Among the 15 analyzable pairs, 12 were scored as genetically homologous. For four out of these 12 genetically homologous pairs also tested by WGS, the very high genetic relatedness of D_0_ and D_relapse_ isolates was confirmed. Contrary to most of *P. vivax* relapses studied in Thailand where heterologous relapses are the norm [45] (exception made of the relapses observed in young infants that were mainly homologous [46]), we report high percentage of homologous relapses. This has to be interpreted in the light of two key features of our study: (i) we studied imported malaria cases; therefore patients mainly live in metropolitan France with short travel to Africa limiting the probabilities, compared to patients living in endemic area, of a previous *P. ovale wallikeri* infection or multiple infections (two ways to get heterologous relapses [47,48]); and (ii) according to the microsatellite results reported here, the polyclonal infections (second way to get heterologous relapses [45]) rate in *P. ovale wallikeri* isolates from imported malaria appears to be much less frequent compared to the rate reported in *P. vivax* (10% of polyclonal infection for *P. ovale wallikeri* in our work *vs* a mean proportion of polyclonal infections of 43% for *P. vivax* in endemic zone [49]).

The genetic relatedness within the three last pairs (14, 15 and 17; Table S7) was more complex. For pairs 14 and 17, D_0_ isolates were polyclonal while D_relapse_ carried no new alleles, confirming their high genetic relatedness. Interestingly, D_relapse_ from pair 14 was monoclonal contrary to relapse from pair 17 that was polyclonal. This may result from different ways of hypnozoite reactivation, stochastic for pair 14 (*i*.*e*. one hypnozoite reactivation) and external stimulus for pair 17 (*i*.*e*. at least two hypnozoites reactivation)[47]. Finally, the two isolates from pair 15 were both monoclonal and differed at a single marker by just one tandem repeat. Although initially classified as heterologous, the new allele in D_relapse_ could be simply derived by mutation of the D_0_ allele early during the course of the relapse infection. Slippage during the PCR was excluded since PCR was repeated.

In conclusion, using a novel set of eight microsatellite markers we genotyped for the first time *P. ovale wallikeri* isolates from primary and relapse infections in the context of imported malaria and low multiplicity of infections. As expected, most paired primary and relapse isolates appeared to be highly genetically related. WGS should help in the future to get a more complete genetic comparison of D_0_-D_relapse_ isolate pairs and also to look for putative genetic markers associated with *P. ovale wallikeri* dormancy.

## Declarations Data availability

The genome Illumina sequencing reads from the *P. ovale wallikeri* samples were deposited in the European Nucleotide Archive under the accession number PRJEB51041.

## Authors’ contributions

VJ, SH and JC designed the study. VJ and ECI performed the experimentations. VJ, ECI, EG and RC analyzed the data. VJ, EG and RC performed the bioinformatic analyses. VJ wrote the article and all authors reviewed the manuscript. SH supervised the study. There is no conflict of interests to disclose.

## Acknowledgments

We thank all collaborating individuals for their participation in collecting materials and data (referred as the French National Reference Center for Imported Malaria Study Group), listed in alphabetical order (last name): Aboubacar Ahmed, Angebault Cécile, Angoulvant Adela, Argy Nicolas, Azjenberg Daniel, Belkadi Ghania, Bellanger Anne-Pauline, Bemba Dieudonné, Blaize Marion, Botterel Françoise, Bougnoux Marie-Elisabeth, Brun Sophie, Buret Bernadette, Chevrier Sylviane, Clauser Sylvain, Dahane Naima, Dannaoui Eric, Dard Cécile, Dardé Marie-Laure, de Gentile Ludovic, de Suremain Nathalie, Debourgogne Anne, Delaval Anne, Deleplancque Anne-Sophie, Desoubeaux Guillaume, Durand Rémy, Durieux Marie-Fleur, Dutoit Emmanuel, Eloy Odile, Fenneteau Odile, Gargala Gilles, Godineau Nadine, Guennouni Nadia, Guinard Jérôme, Hamane Samia, Herault Etienne, Larreché Sébastien, Lavergne Rose-Anne, Marteau Anthony, Mazars Edith, Moreno-Sabater Alicia, Morio Florian, Nourrisson Céline, Perraud-Cateau Estelle, Pons Denis, Pull Lauren, Quinio Dorothée, Raffenot Didier, Silva Muriel, Thellier Marc, Tielli Alexandra, Toubas Dominique.

