## Supplemental material for "Genetic profiling of *Plasmodium ovale wallikeri* relapses with microsatellite markers and whole-genome sequencing"

### Appendix

#### I. Supplemental figures

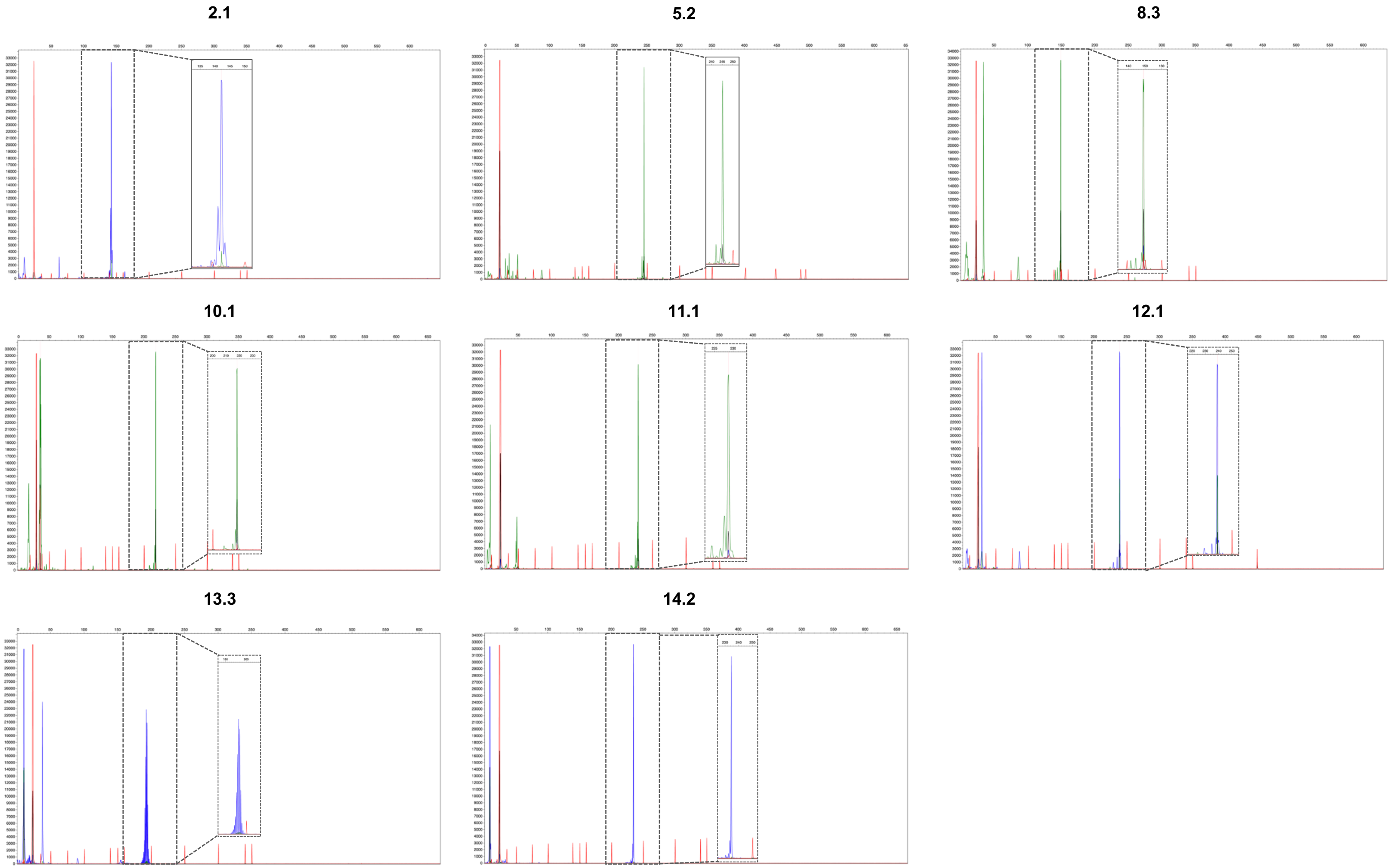

**Figure S1 – Microsatellite profile examples for the eight microsatellites loci with the GeneScan$^{TM}$ 350 or 500 ROX$^{TM}$ size markers.** Electrophoregrams were visualized with the Microsatellite Analysis software on the Thermo Fisher Cloud.

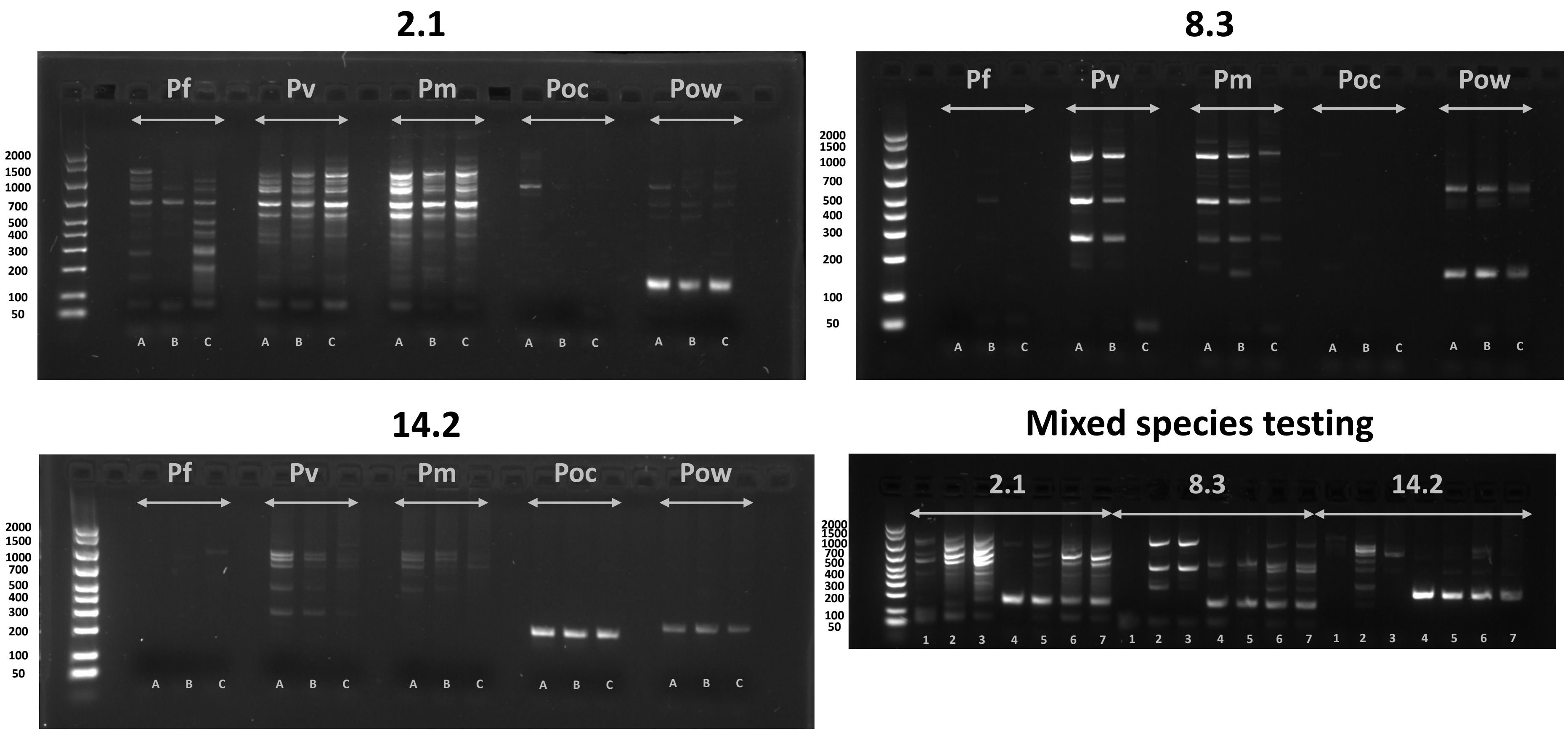

**Figure S2 – 2% gel agarose stained with 10000X GelRed® for markers 2.1, 8.3 and 14.2 and mixed species testing.**
A, B and C correspond to 62-57, 63-58 and 64-59°C annealing temperatures (of the first and second cycle) of the touchdown PCRs. Pf stands for *P. falciparum* (parasite density = 12,000 p/µL), Pv for *P. vivax* (9,000 p/µL), Pm for *P. malariae* (9,000 p/µL), Poc for *P. ovale curtisi* and Pow for *P. ovale wallikeri* (4,500 p/µL). 1 to 7 correspond respectively to Pf alone, Pm alone, Pv alone, Pow alone, Pf + Pow, Pm + Pow and Pv + Pow.
Amplisize molecular ruler (Bio-Rad) was used as DNA size marker and is indicated on the left side of each gel.

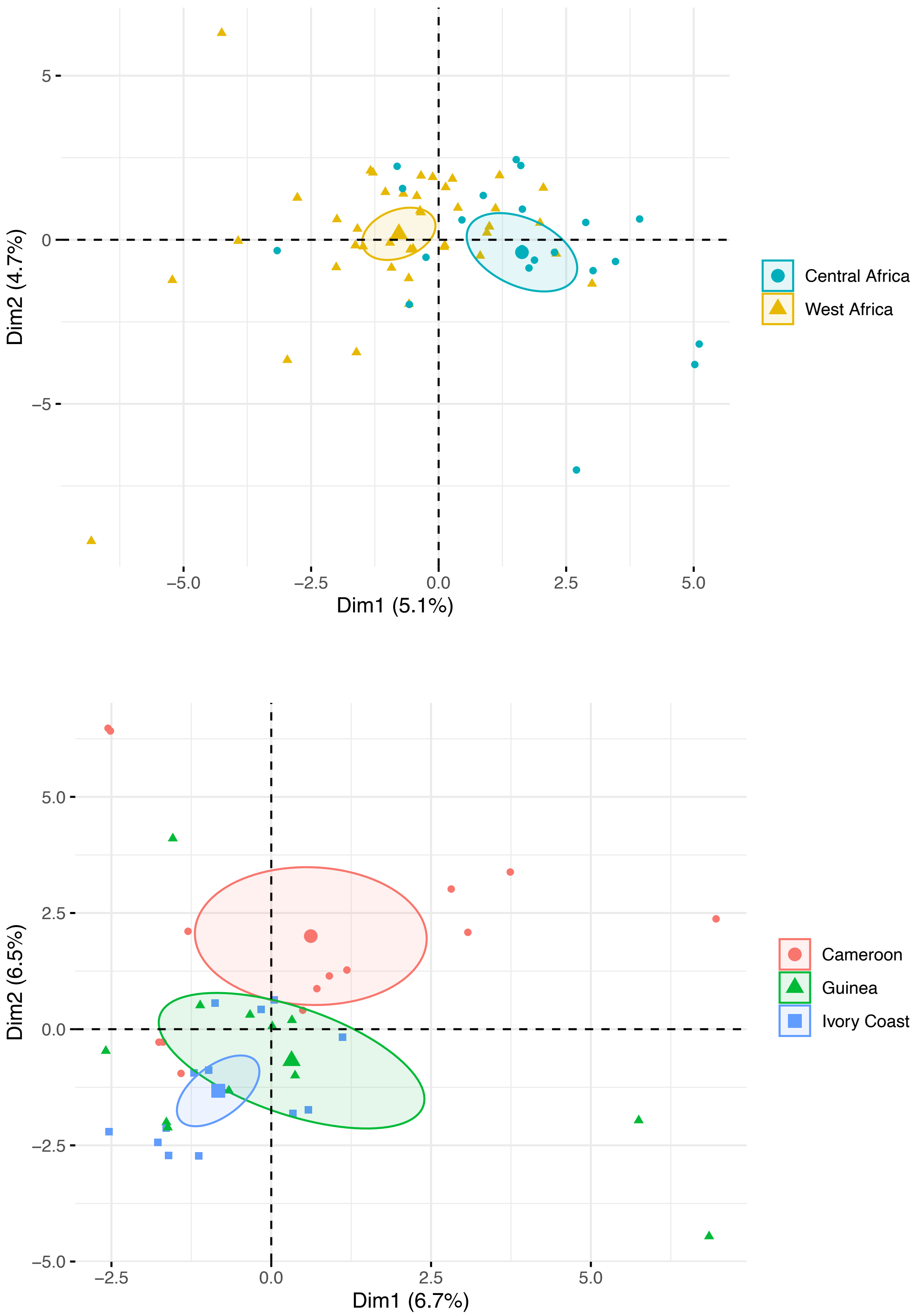

**Figure S3 – Principal Component Analysis based on the microsatellite allele sizes for the eight markers for isolates from A) West (brown) or Central Africa (blue) or from B) Cameroon (red), Guinea (green) or Ivory Coast (blue).**

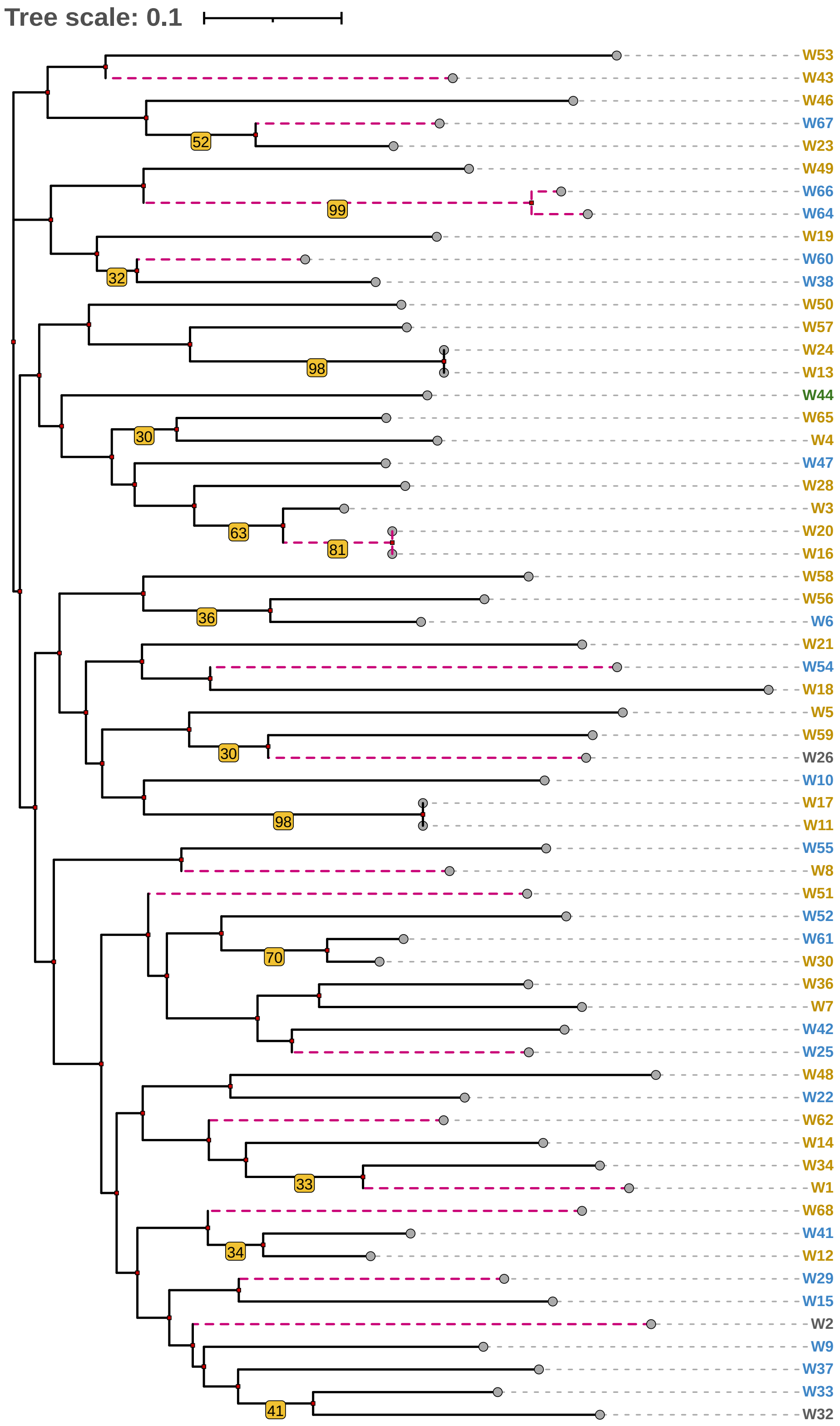

**Figure S4 - Phylogenetic tree built using the neighbor-joining method based on the microsatellite allele sizes of *P. ovale wallikeri* isolates with a complete eight-marker haplotype**.

Isolates are classified as from West Africa (brown), Central Africa (blue), East Africa (green) and multiples countries (gray). Branches that support isolates from relapse pairs are colored in purple. Bootstrap values > 30 are indicated.

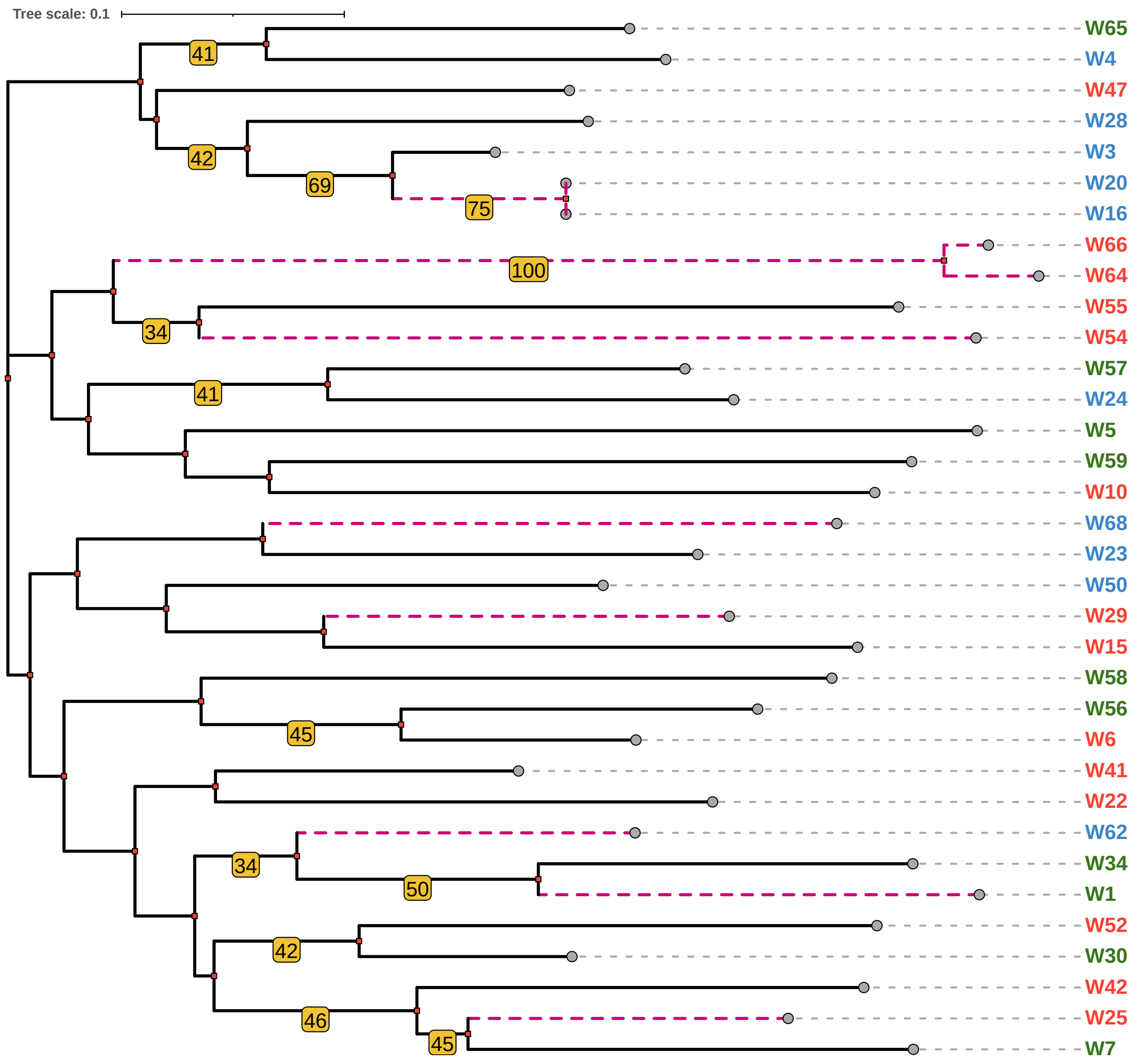

**Figure S5 - Phylogenetic tree built using the neighbor-joining method based on the microsatellite allele sizes of *P. ovale wallikeri* isolates from Cameroon (in red), Guinea (in green) and Ivory Coast (in blue) with a complete eight-marker haplotype.**
Branches that support isolates from relapse pairs are colored in purple. Bootstrap values > 30 are indicated.

#### II. Supplemental tables

**Table S1 – Epidemiological, biological and clinical characteristics of the $D_0$ and $D_{relapse}$ infections**

| Characteristics | $D_0$ (n=17) | $D_{relapse}$ (n=17) |
|---|---|---|
| Age (years), median [IQR] | 29 [17-34] | |
| Gender (% female) | 35.2 | |
| Country of contamination, N | | |
| Cameroon | 4 | |
| Central African Republic | 1 | |
| Republic of Congo | 1 | |
| Guinea | 1 | |
| Ivory Coast | 5 | |
| Nigeria | 1 | |
| Senegal | 1 | |
| Togo | 1 | |
| Several countries | 2 | |
| Country of living, N | | |
| Metropolitan France | 15 | |
| Burkina-Faso | 1 | |
| Missing data | 1 | |
| Duration of travel (days), median [IQR] | 30 [22-52] | |
| Latency period (days), median [IQR] | 50 [22-77] | |
| Relapse delay (days), median [IQR] | 108 [60-145] | |
| Parasite density (p/µL), median [IQR] | 1,350 [720-2,250] | 788 [228-1,861] |
| Leucocytes count (G/L), median [IQR] | 6 [3.9-7.4] | 5.2 [3.8-6.1] |
| Platelets count (G/L), median [IQR] | 103 [66-122] | 111 [76-133] |
| Hemoglobin (g/L), median [IQR] | 130 [116-138] | 125 [120-139] |
| Chemoprophylaxis: N (%) | | |
| Yes | 8 (47) | |
| Complete | 4 (50) | |
| Incomplete | 4 (50) | |
| No | 7 (41) | |
| Missing data | 2 (12) | |
| Clinical categorization, N (%) | | |
| Uncomplicated malaria, | 17 (100) | 11 (100) |
| Missing data | 0 | 6 |
| Treatment, N (%) | | |
| Chloroquine | 8 (57) | 5 (45.5) |
| Atovaquone-proguanil | 3 (21.5) | 2 (18) |
| ACT | 3 (21.5) | 3 (27.5) |
| Quinine | 0 (0) | 1 (9) |
| Missing data | 3 | 6 |

IQR: InterQuartil Interval; ACT: Artemisinin-Combination Therapy; Latency period is the delay between the return from endemic country and the symptom onset

**Table S2 – Detailed epidemiological data for the seventeen pairs of relapsing isolates.**

| Relapse | Country* | Residency | Length of stay | Relapse delay+ | Negative follow-up Parasitaemia | Treatment of the 1st infection$ | Radical treatment 1st infection | Radical treatment 2nd infection |
|---|---|---|---|---|---|---|---|---|
| Pair 1 | Ivory Coast | France | 28 | 122 | D3 | AP | No | Yes |
| Pair 2 | Cameroon | France | 387 | 33 | None | CQ | No | No |
| Pair 3 | Cameroon | France | 17 | 345 | D3, D7, D28 | CQ | No | NR |
| Pair 4 | Senegal | France | NR | 134 | None | AL | No | NR |
| Pair 5 | Cameroon | France | 11 | 28 | D7 | AL | No | NR |
| Pair 6 | Togo | France | 24 | 153 | D3 | AL | No | Yes |
| Pair 7 | Ivory Coast | France | 32 | 61 | None | CQ | No | NR |
| Pair 8 | Guinea | France | 60 | 78 | D3, D5 | AP | No | No |
| Pair 9 | Several | France | 31 | 120 | D3, D7, D28 | CQ | NR | NR |
| Pair 10 | Nigeria | France | 23 | 53 | None | NA | NR | Yes |
| Pair 11 | Several | BF | 694 | 39 | D3 | CQ | No | Yes – complete |
| Pair 12 | Ivory Coast | NR | NR | 257 | D7, D21, D28 | NA | No – G6PD deficiency | NR |
| Pair 13 | Ivory Coast | France | 22 | 60 | D28 | CQ | No | Yes – complete |
| Pair 14 | Ivory Coast | France | 33 | 145 | D5 | DHA-PPQ | No – G6PD deficiency | No – G6PD deficiency |
| Pair 15 | Cameroon | France | 21 | 62 | D7 | AP | No – G6PD deficiency | No – G6PD deficiency |
| Pair 16 | Congo | France | NR | 189 | D3, D7, D28 | CQ | No – G6PD deficiency | No – G6PD deficiency |
| Pair 17 | CAR | France | 31 | 108 | D4, D7, D28 | NA | No – G6PD deficiency | No – G6PD deficiency |

*CAR stands for Central African Republic, BF for Burkina Faso, NR for Not Reported.

+Delay between the $D_0$ of the primary infection and the $D_0$ of the second infection

$AP stands for Atovaquone-Proguanil, CQ for Chloroquine, AL for Artemeter-Lumefantrine and DHA-PPQ for Dihydroartemisinin-Piperaquine

**Table S3 – Microsatellite allele length for the eight markers for the 69 *P. ovale wallikeri* isolates.**
Relapse isolate with identical haplotype than primary isolate were not included.

| Isolate | Country | 2.1 | 5.2 | 8.3 | 10.1 | 11.1 | 12.1 | 13.3 | 14.2 | Haplotype |
|---|---|---|---|---|---|---|---|---|---|---|
| W1* | Guinea | 149 | 250 | 127 | 203 | 185 | 235 | 192 | 239 | H1 |
| W2* | Several countries | 149 | 253 | 144 | 219 | 246 | 267 | 195 | 266 | H2 |
| W3 | Ivory Coast | 142 | 243 | 149 | 219 | 207 | 239 | 189 | 235 | H3 |
| W4 | Ivory Coast | 142 | 243 | 166 | 219 | 221 | 239 | 189 | 244 | H4 |
| W5 | Guinea | 146 | 257 | 149 | 212 | 197 | 248 | 189 | 278 | H5 |
| W6 | Cameroon | 149 | 250 | 144 | 222 | 207 | 259 | 189 | 231 | H6 |
| W7 | Guinea | 152 | 250 | 139 | 219 | 211 | 235 | 267 | 252 | H7 |
| W8* | Nigeria | 142 | 248 | 144 | 222 | 215 | 235 | 210 | 239 | H8 |
| W9 | Centrafrique | 149 | 243 | 139 | 219 | 197 | 243 | 195 | 239 | H9 |
| W10 | Cameroon | 139 | 263 | 144 | 222 | 202 | 248 | 216 | 231 | H10 |
| W11 | Togo | 139 | 243 | 144 | 222 | 215 | 252 | 189 | 235 | H11 |
| W12 | Mali | 149 | 243 | 139 | 222 | 207 | 239 | 189 | 231 | H12 |
| W13 | Burkina Faso | 142 | 237 | 149 | 222 | 229 | 235 | 189 | 278 | H13 |
| W14 | Benin | 149 | 237 | 127 | 222 | 207 | 235 | 192 | 282 | H14 |
| W15 | Cameroon | 149 | 243 | 241, 245 | 222 | 202 | 252 | 195 | 227 | H15 |
| W16* | Ivory Coast | 142 | 243 | 149 | 219 | 229 | 227, 239 | 189 | 235 | H16a |
| W17 | Togo | 139 | 243 | 144 | 222 | 215 | 252 | 189 | 235 | H11 |
| W18 | Ghana | 136 | 237 | 473 | 232 | 233 | 256 | 195 | 344 | H17 |
| W19 | Togo | 142 | 250 | 144 | 239 | 207 | 239 | 189 | 274 | H18 |
| W20* | Ivory Coast | 142 | 243 | 149 | 219 | 229 | 239 | 189 | 235 | H16b |
| W21 | Sierra Leone | 132 | 243 | 135 | 210 | 233 | 248 | 192 | 231 | H19 |
| W22 | Cameroon | 149 | 233 | 144 | 222 | 193 | 235 | 189 | 235 | H20 |
| W23 | Ivory Coast | 142 | 243 | 139 | 206 | 207 | 239 | 207 | 231 | H21 |
| W24 | Ivory Coast | 142 | 237 | 149 | 222 | 229 | 235 | 189 | 278 | H13 |
| W25* | Cameroon | 152 | 243 | 139 | 219 | 207 | 235 | 207 | 256 | H22 |

| | | | | | | | | | | |
|---|---|---|---|---|---|---|---|---|---|---|
| W26* | Several countries | 156 | 245 | 149 | 222 | 229 | 250 | 195 | 278 | H23 |
| W27 | Guinea | 142 | 243 | 154 | 219 | 207 | nd | 189 | 256 | H24 |
| W28 | Ivory Coast | 136, 142 | 245 | 154 | 219 | 211 | 235 | 189 | 235 | H25 |
| W29* | Cameroon | 149 | 243 | 139 | 222 | 200 | 267 | 207 | 231 | H26 |
| W30 | Guinea | 149 | 243 | 144 | 219 | 207 | 235 | 189 | 244 | H27 |
| W31 | Guinea | 149 | 245 | nd | nd | 233 | 239 | 204 | 248 | H28 |
| W32 | Several countries | 149 | 253 | 158 | 219 | 221 | 248 | 195 | 231 | H29 |
| W33 | Congo | 149 | 243 | 158 | 219 | 197 | 235 | 195 | 231 | H30 |
| W34 | Guinea | 149 | 250 | 127 | 206 | 207 | 235 | 204 | 235 | H31 |
| W35 | Niger | 149 | nd | 144 | 219 | 221 | 352, 355 | 189 | 256 | H32 |
| W36 | Senegal | 149 | 250 | 139 | 219 | 211 | 239 | 228 | 244 | H33 |
| W37 | Congo | 149 | 243 | 154 | 219 | 211 | 235 | 198 | 227 | H34 |
| W38 | Gabon | 142 | 245 | 127 | 219 | 207 | 239 | 189 | 259 | H35 |
| W39 | Ivory Coast | 142 | 245 | 135 | nd | 233 | nd | nd | 266 | H36 |
| W40 | Ivory Coast | 142 | 243 | 149 | 219 | 229 | nd | 189 | 235 | NC |
| W41 | Cameroon | 149 | 243 | 139 | 222 | 225 | 235 | 189 | 231 | H37 |
| W42 | Cameroon | 149 | 250 | 139 | 219 | 193 | 235 | 207 | 266 | H38 |
| W43* | Senegal | 142 | 237 | 139 | 219 | 197 | 235 | 213 | 252 | H39 |
| W44 | Rwanda | 142 | 245 | 144 | 225 | 200 | 227 | 189 | 244 | H40 |
| W45 | Ivory Coast | nd | 243 | 144 | 222 | 225 | nd | nd | 248 | H41 |
| W46 | Nigeria | 142 | 237 | 503 | 203 | 207 | 252 | 204 | 231 | H42 |
| W47 | Cameroon | 142 | 243 | 139 | 219 | 211 | 270 | 189 | 231 | H43 |
| W48 | Ghana | 150 | 233 | 170 | 222 | 250 | 259 | 198 | 235 | H44 |
| W49 | Sierra Leone | 142 | 250 | 139 | 206 | 185 | 235 | 192 | 278 | H45 |
| W50 | Ivory Coast | 142 | 243 | 166 | 222 | 202 | 235 | 192 | 231 | H46 |
| W51* | Togo | 149 | 237 | 139 | 222 | 207 | 248 | 267 | 244 | H47 |
| W52 | Cameroon | 158 | 250 | 357 | 219 | 207 | 235 | 189 | 244 | H48 |
| W53 | Nigeria | 142 | 233 | 618 | 212 | 193 | 243 | 210 | 244 | H49 |

| | | | | | | | | | | |
|---|---|---|---|---|---|---|---|---|---|---|
| W54* | Cameroon | 142 | 243 | 464 | 222 | 237 | 256 | 222 | 259 | H50 |
| W55 | Cameroon | 146 | 250 | 135 | 222 | 215 | 239 | 222 | 248 | H51 |
| W56 | Guinea | 149 | 250 | 144 | 232 | 207 | 252 | 189 | 235 | H52 |
| W57 | Guinea | 142 | 237 | 149 | 222 | 207 | 235 | 183 | 231 | H53 |
| W58 | Guinea | 142 | 253 | 139 | 232 | 189 | 256 | 189 | 231 | H54 |
| W59 | Guinea | 136 | 243 | 144 | 222 | 221 | 248 | 243 | 278 | H55 |
| W60* | Congo | 142 | 245 | 139 | 219 | 207 | 239 | 192 | 227 | H56 |
| W61 | Gabon | 149 | 243 | 144 | 219 | 207 | 243 | 189 | 244 | H57 |
| W62* | Ivory Coast | 149 | 243 | 127 | 219 | 202 | 235 | 189 | 235 | H58 |
| W63 | Ivory Coast | 142 | 253 | 144 | 222 | 207 | 250 | nd | 239 | H59 |
| W64* | Cameroon | 142 | 253 | 158 | 203 | 229 | 243 | 192 | 266 | H60 |
| W65 | Guinea | 142 | 243 | 139 | 219 | 250 | 250 | 189 | 244 | H61 |
| W66 | Cameroon | 142 | 250 | 158 | 203 | 229 | 243 | 192 | 266 | H62 |
| W67* | Central African Republic | 142, 146 | 243 | 139 | 206 | 215, 225 | 239, 259 | 204 | 231, 235 | H63 |
| W68* | Ivory Coast | 149, 156 | 243, 245 | 135, 139 | 215, 219 | 207, 250 | 235, 239 | 213 | 231 | H64 |
| W69 | Ivory Coast | 142 | 237 | 149 | 222 | 279 | 267 | nd | 282 | H65 |

NC: Not Classifiable; nd: not determined, nt: not tested. Several countries correspond to patient that through more than one country (in West Africa) during their journey.
*Primary isolate from relapse infection

**Table S4 – Allele frequencies and sample sizes (n) for the eight microsatellite markers from the total cohort and 3 populations of *P. ovale wallikeri*.**

| Allele Length (bp) | Africa | | | |
|---|---|---|---|---|
| | Total cohort | Cameroon | Guinea | Ivory Coast |
| **2.1** | | | | |
| 132 | 0.014 | / | / | / |
| 136 | 0.042 | / | 0.083 | 0.063 |
| 139 | 0.042 | 0.071 | / | / |
| 142 | 0.437 | 0.286 | 0.333 | 0.75 |
| 146 | 0.042 | 0.071 | 0.083 | / |
| 149 | 0.338 | 0.429 | 0.417 | 0.125 |
| 150 | 0.014 | / | / | / |
| 152 | 0.028 | 0.071 | 0.083 | / |
| 156 | 0.028 | / | / | 0.063 |
| 159 | 0.014 | 0.071 | / | / |
| n | 71 | 14 | 12 | 16 |
| **5.2** | | | | |
| 233 | 0.043 | 0.071 | / | / |
| 237 | 0.13 | / | 0.083 | 0.125 |
| 243 | 0.420 | 0.429 | 0.333 | 0.625 |
| 245 | 0.116 | / | 0.083 | 0.188 |
| 248 | 0.014 | / | / | / |
| 250 | 0.174 | 0.357 | 0.333 | / |
| 253 | 0.072 | 0.071 | 0.083 | 0.063 |
| 257 | 0.014 | / | 0.083 | / |
| 263 | 0.014 | 0.071 | / | / |
| n | 69 | 14 | 12 | 16 |
| **8.3** | | | | |
| 127 | 0,070 | / | 0.182 | 0.059 |
| 135 | 0,056 | 0.067 | / | 0.118 |
| 139 | 0,268 | 0.333 | 0.273 | 0.176 |
| 144 | 0,225 | 0.200 | 0.273 | 0.118 |
| 149 | 0,141 | / | 0.182 | 0.353 |
| 154 | 0,0423 | / | 0.091 | 0.059 |
| 158 | 0,056 | 0.133 | / | / |
| 166 | 0,0282 | / | / | 0.118 |
| 170 | 0,0141 | / | / | / |
| 241 | 0,0141 | 0.067 | / | / |
| 245 | 0,0141 | 0.067 | / | / |
| 357 | 0,0141 | 0.067 | / | / |
| 464 | 0,0141 | 0.067 | / | / |
| 473 | 0,0141 | / | / | / |
| 503 | 0,0141 | / | / | / |
| 518 | 0,0141 | / | / | / |
| n | 71 | 15 | 11 | 17 |
| **10.1** | | | | |
| 203 | 0,059 | 0.143 | 0.091 | / |
| 206 | 0,059 | / | 0.091 | 0.067 |
| 210 | 0,015 | / | / | / |
| 212 | 0,029 | / | 0.091 | / |
| 215 | 0,015 | / | / | 0.067 |
| 219 | 0,397 | 0.286 | 0.364 | 0.533 |
| 222 | 0,353 | 0.571 | 0.182 | 0.333 |

| | | | | |
|---|---|---|---|---|
| 225 | 0,015 | / | / | / |
| 232 | 0,044 | / | 0.182 | / |
| 239 | 0,015 | / | / | / |
| n | 68 | 14 | 11 | 15 |
| **11.1** | | | | |
| 185 | 0,028 | / | 0.083 | / |
| 189 | 0,014 | / | 0.083 | / |
| 193 | 0,042 | 0.143 | / | / |
| 197 | 0,056 | / | 0.083 | / |
| 200 | 0,028 | 0.071 | / | / |
| 202 | 0,056 | 0.143 | / | 0.125 |
| 207 | 0,282 | 0.214 | 0.417 | 0.25 |
| 211 | 0,070 | 0.071 | 0.083 | 0.063 |
| 215 | 0,070 | 0.071 | / | / |
| 221 | 0,056 | / | 0.083 | 0.063 |
| 225 | 0,042 | 0.071 | / | 0.063 |
| 229 | 0,113 | 0.143 | / | 0.25 |
| 233 | 0,056 | / | 0.083 | 0.063 |
| 237 | 0,014 | 0.071 | / | / |
| 246 | 0,014 | / | / | / |
| 250 | 0,042 | / | 0.083 | 0.063 |
| 279 | 0,014 | / | / | 0.063 |
| n | 71 | 14 | 12 | 16 |
| **12.1** | | | | |
| 227 | 0,029 | / | / | 0.071 |
| 235 | 0,319 | 0.357 | 0.455 | 0.357 |
| 239 | 0,203 | 0.071 | 0.091 | 0.429 |
| 243 | 0,072 | 0.143 | / | / |
| 248 | 0,087 | 0.071 | 0.182 | / |
| 250 | 0,043 | / | 0.091 | 0.071 |
| 252 | 0,072 | 0.071 | 0.091 | / |
| 256 | 0,043 | 0.071 | 0.091 | / |
| 259 | 0,043 | 0.071 | / | / |
| 267 | 0,043 | 0.071 | / | 0.071 |
| 270 | 0,014 | 0.071 | / | / |
| 352 | 0,014 | / | / | / |
| 356 | 0,014 | / | / | / |
| n | 69 | 14 | 11 | 14 |
| **13.3** | | | | |
| 183 | 0,015 | / | 0.083 | / |
| 189 | 0,431 | 0.357 | 0.5 | 0.727 |
| 192 | 0,123 | 0.143 | 0.083 | 0.091 |
| 195 | 0,108 | 0.071 | / | / |
| 198 | 0,031 | / | / | / |
| 204 | 0,062 | / | 0.167 | / |
| 207 | 0,062 | 0.214 | / | 0.091 |
| 210 | 0,031 | / | / | / |
| 213 | 0,031 | / | / | 0.091 |
| 216 | 0,015 | 0.071 | / | / |
| 222 | 0,031 | 0.143 | / | / |
| 228 | 0,015 | / | / | / |
| 243 | 0,015 | / | 0.083 | / |
| 267 | 0,031 | / | 0.083 | / |
| n | 65 | 14 | 12 | 11 |
| 14.2 | | | | |

| | | | | |
|---|---|---|---|---|
| 227 | 0,043 | 0.071 | / | / |
| 231 | 0,229 | 0.357 | 0.167 | 0.200 |
| 235 | 0,186 | 0.071 | 0.167 | 0.400 |
| 239 | 0,057 | / | 0.083 | 0.067 |
| 244 | 0,129 | 0.071 | 0.167 | 0.067 |
| 248 | 0,043 | 0.071 | 0.083 | 0.067 |
| 252 | 0,029 | / | 0.083 | / |
| 256 | 0,043 | 0.071 | 0.083 | / |
| 259 | 0,029 | 0.071 | / | / |
| 266 | 0,071 | 0.214 | / | 0.067 |
| 274 | 0,014 | / | / | / |
| 278 | 0,086 | / | 0.167 | 0.067 |
| 282 | 0,029 | / | / | 0.067 |
| 344 | 0,014 | / | / | / |
| n | 70 | 14 | 12 | 15 |

**Table S5 – Combined probabilities of the microsatellite markers.**

Markers are classified in ascending order of probability for each marker. Combined probabilities value is the probability to have the exact same haplotype for two random isolates.

| Microsatellite marker | Probability per marker ($p=\sum p_i^2$) | Combined probabilities ($\pi P_i$) | Combined probabilities values |
|---|---|---|---|
| **11.1** | 0.122 | $P1= P_{11.1}$ | 0.122 |
| **14.2** | 0.127 | $P2= P1 \times P_{14.2}$ | $1.5 \times 10^{-2}$ |
| **8.3** | 0.158 | $P3=P2 \times P_{8.3}$ | $2.5 \times 10^{-3}$ |
| **12.1** | 0.170 | $P4=P3 \times P_{12.1}$ | $4.2 \times 10^{-4}$ |
| **13.3** | 0.226 | $P5=P4 \times P_{13.3}$ | $9.4 \times 10^{-5}$ |
| **5.2** | 0.245 | $P6=P5 \times P_{5.2}$ | $2.3 \times 10^{-5}$ |
| **10.1** | 0.293 | $P7=P6 \times P_{10.1}$ | $6.8 \times 10^{-6}$ |
| **2.1** | 0.312 | $P8=P7 \times P_{2.1}$ | $2.1 \times 10^{-6}$ |

**Table S6 – Patterns of *Plasmodium ovale wallikeri* diversity.**

| Population | N | $H_E$ +/- SD | A +/- SD |
|---|---|---|---|
| West Africa | | | |
| Guinea | 11.6 +/- 0.5 | 0.829 +/- 0.06 | 6.25 +/- 1.16 |
| Ivory Coast | 15 +/- 1.9 | 0.682 +/- 0.168 | 5.5 +/- 1.93 |
| Central Africa | | | |
| Cameroon | 14.1 +/- 0.4 | 0.810 +/- 0.102 | 6.75 +/- 2.12 |

N: number of isolates, averaged over all markers; $H_E$: expected heterozygosity averaged all markers; A: mean number of alleles per marker.
SD stands for Standard Deviation.

**Table S7 – Microsatellite allele sizes obtained for $D_0$ and $D_{relapse}$ isolate pairs.**
Parasite density in p/µL is also indicated.

| Relapse | Isolate[$] | Episode | Parasite density | 2.1 | 5.2 | 8.3 | 10.1 | 11.1 | 12.1 | 13.3 | 14.2 | Conclusion |
|---|---|---|---|---|---|---|---|---|---|---|---|---|
| Pair 1* | W20 | Primary | 1,350 | 142 | 243 | 149 | 219 | 229 | 239 | 190 | 235 | Homologous |
| | W20bis | Relapse | 775 | 142 | 243 | 149 | 219 | 229 | 239 | 190 | 235 | |
| Pair 2* | W25 | Primary | 1800 | 152 | 243 | 139 | 219 | 207 | 235 | 206 | 256 | Homologous |
| | W25bis | Relapse | 450 | 152 | 243 | 139 | 219 | 207 | 235 | 206 | 256 | |
| Pair 3 | W29 | Primary | 450 | 149 | 243 | 139 | 222 | 200 | 267 | 206 | 231 | Homologous |
| | W29bis | Relapse | Not recorded | 149 | 243 | 139 | 222 | 200 | 267 | 206 | 231 | |
| Pair 4* | W43 | Primary | 14,850 | 142 | 237 | 139 | 219 | 197 | 235 | 214 | 252 | Homologous |
| | W43bis | Relapse | 22,500 | 142 | 237 | 139 | 219 | 197 | 235 | 214 | 252 | |
| Pair 5* | W54 | Primary | 23,850 | 142 | 243 | 464 | 222 | 237 | 256 | 223 | 259 | Homologous |
| | W54bis | Relapse | 248 | 142 | 243 | 464 | 222 | 237 | 256 | 223 | 259 | |
| Pair 6 | W51 | Primary | 720 | 149 | 237 | 139 | 222 | 207 | 248 | 267 | 244 | NC |
| | W51bis | Relapse | 153 | 149 | Failed | Failed | Failed | Failed | 248 | 267 | Failed | |
| Pair 7 | W62 | Primary | 2,250 | 149 | 243 | 127 | 219 | 202 | 235 | 190 | 235 | Homologous |
| | W62bis | Relapse | 72 | 149 | 243 | 127 | 219 | 202 | 235 | 190 | 235 | |
| Pair 8 | W1 | Primary | 9,300 | 149 | 250 | 127 | 203 | 185 | 235 | 193 | 239 | Homologous |
| | W1bis | Relapse | 13,500 | 149 | 250 | 127 | 203 | 185 | 235 | 193 | 239 | |
| Pair 9 | W2 | Primary | 1,350 | 149 | 253 | 144 | 219 | 246 | 267 | 195 | 266 | Homologous |
| | W2bis | Relapse | 9,900 | 149 | 253 | 144 | 219 | 247 | 267 | 195 | 266 | |
| Pair 10 | W8 | Primary | 1,000 | 142 | 248 | 144 | 222 | 215 | 235 | 210 | 239 | Homologous |
| | W8bis | Relapse | 1,800 | 142 | 248 | 144 | 222 | 215 | 235 | 210 | 239 | |
| Pair 11 | W26 | Primary | 720 | 156 | 245 | 149 | 222 | 229 | 250 | 195 | 278 | Homologous |
| | W26bis | Relapse | 1,922 | 156 | 245 | 149 | 222 | 229 | 250 | 195 | 278 | |
| Pair 12 | W16 | Primary | 1,620 | 142 | 243 | 149 | 219 | 239 | 239 | 190 | 235 | Homologous |
| | W16bis | Relapse | 1,350 | 142 | 243 | 149 | 219 | 239 | 239 | 190 | 235 | |
| Pair 13 | Not included | Primary | 4,760 | 142 | Failed | 139 | Failed | 197 | NR | NR | NR | NC |

| | | | | | | | | | | | | |
|---|---|---|---|---|---|---|---|---|---|---|---|---|
| | Not included | Relapse | 81 | Failed | 248 | 139 | Failed | Failed | NR | NR | NR | |
| Pair 14 | W68 | Primary | 640 | 149, 156 | 243, 245 | 135, 139 | 215, 219 | 207, 250 | 235, 239 | 213 | 231 | Related |
| | W68bis | Relapse | 1,840 | 156 | 243 | 139 | 215 | 250 | 235 | 213 | 231 | |
| Pair 15 | W64 | Primary | 1,320 | 142 | 253 | 158 | 203 | 229 | 243 | 193 | 266 | Heterologous |
| | W66 | Relapse | 168 | 142 | 250 | 158 | 203 | 229 | 243 | 193 | 266 | |
| Pair 16 | W60 | Primary | 320 | 142 | 245 | 139 | 219 | 207 | 239 | 193 | 227 | Homologous |
| | W60bis | Relapse | 800 | 142 | 245 | 139 | 219 | 207 | 239 | 193 | 227 | |
| Pair 17 | W67 | Primary | 60 | 142, 146 | 243 | 139 | 206 | 215, 225 | 239, 259 | 203 | 231, 235 | Related |
| | W67bis | Relapse | 450 | 142, 146 | 243 | 139 | 206 | 225 | 259 | 203 | 231 | |

$Refers to the isolate name from Table S2. bis refers to the relapse isolate identical to the primary isolate and not included in the genetic diversity analysis.
*Pairs for which NGS was performed.
NC stands for Not Classifiable, NR for Not Realized (not enough DNA).

**Table S8 – Number of genome-wide SNPs before and after filtration.**
Percentage of reads that mapped to reference genome (GCA_900090025.2) and percentage of the genome covered with at least 5 reads are also indicated.

| Pair (isolate[&]) | ENA | Episode | Reads (M) | % mapped | 5X coverage | Total SNPs* | Chr SNPs | Hz SNPs[$] | Het SNPs[£] |
|---|---|---|---|---|---|---|---|---|---|
| Relapse pair 1 (W20-W20bis) | ERS14392655 | Primary | 27 | 31 | 74% | 4,173 | 2,015 | 1,956 | 59 |
| | ERS14392659 | Relapse | 24 | 27 | 61% | 2,092 | 1,119 | 1,088 | 31 |
| Relapse pair 2 (W25-W25bis) | ERS14392656 | Primary | 32 | 38 | 86% | 8,511 | 2,602 | 2,436 | 166 |
| | ERS14392660 | Relapse | 12 | 75 | 73% | 5,062 | 2,967 | 2,655 | 312 |
| Relapse pair 4 (W43-W43bis) | ERS14392657 | Primary | 27 | 97 | 92% | 16,236 | 5,564 | 4,818 | 746 |
| | ERS14392661 | Relapse | 23 | 92 | 92% | 15,191 | 5,422 | 4,736 | 686 |
| Relapse pair 5 (W54-W54bis) | ERS14392658 | Primary | 31 | 91 | 94% | 16,683 | 5,340 | 4,572 | 768 |
| | ERS14392662 | Relapse | 12 | 96 | 48% | 6,602 | 1,615 | 1,435 | 180 |
| Matched control pair | ERS10659517 | NA | 26 | 95 | 87% | 12,586 | 5,771 | 4,826 | 945 |
| | ERS14392651 | NA | 28 | 95 | 95% | 19,060 | 6,059 | 5,081 | 978 |
| Unmatched control pair 1 | ERS14392655 | NA | 27 | 31 | 74% | 4,173 | 2,015 | 1,956 | 59 |
| | ERS14392654 | NA | 16 | 86 | 95% | 18,210 | 5,453 | 4,942 | 511 |
| Unmatched control pair 2 | ERS14392658 | NA | 31 | 91 | 94% | 16,683 | 5,340 | 4,572 | 768 |
| | ERS10659520 | NA | 31 | 68 | 93% | 11,083 | 4,973 | 4,423 | 550 |

[&]Refers to the isolate name from Table S2.
*SNPs in the chromosomes and in the contigs.
[$]Homozygous SNPs in the chromosomes.
[£]Heterozygous SNPs in the chromosomes.
Chr stands for chromosome and NA for Not Attributable

**Table S9 – Comparison of the SNPs obtained within each pair by NGS.**

| Pair | Isolate[&] | SNPs investigated | Identical SNPs | Wild-Type | % of identical SNPs[$] |
|---|---|---|---|---|---|
| Relapse pair 1 | W20<br>W20bis | 926 | 926 | 0 | 100 |
| Relapse pair 2 | W25<br>W25bis | 1,759 | 1,758 | 1 | 99.94 |
| Relapse pair 4 | W43<br>W43bis | 4,234 | 4,230 | 4 | 99.91 |
| Relapse pair 5 | W54<br>W54bis | 1,319 | 1,302 | 17 | 98.71 |
| Matched control pair | NA<br>NA | 4,284 | 4,281 | 3 | 99.93 |
| Unmatched control pair 1 | W20<br>NA | 2,972 | 941 | 2,031 | 31.66 |
| Unmatched control pair 2 | W54<br>NA | 6,080 | 1,933 | 4,147 | 31.79 |

[&]Refers to the isolate name from Table S2.
[$]Heterozygote SNPs were excluded for the calculation of the percentage.
NA stands for Not Attributable.

#### III. Supplemental methods

Microsatellite development and analysis

The PCR mix contained 0.4 units of GoTaq® Flexi G2 DNA Polymerase (Promega), 200 µM of dNTPs (ThermoFisher Scientific), 1.5 mM of $MgCl_2$ (Promega), 200 nM of forward and reverse primers, 1 µL of DNA and molecular biology grade water to reach a final volume of 10 µL. All PCRs were carried out on an iCycler (BioRad) with the following program: 5 min at 94°C; 10 cycles consisting of 20 sec at 94°C, 30 sec of annealing starting at 62°C (decrease of 0.5°C per cycle) and 25 sec at 68°C; 30 cycles consisting of 20 sec at 94°C, 30 sec at 57°C and 25 sec at 68°C; and a final extension step of 10 min at 68°C.

Selective whole-genome amplification (sWGA)

Each sWGA was performed after methylation digestion of 20 µL of genomic DNA with the McrBC enzyme (New England Biolabs, United Kingdoms) in a mix containing 10 units of McrBC (New England Biolabs), 1× NEBuffer 2 (New England Biolabs), 0.5 µL of 100× BSA (New England Biolabs), 0.5 µl of 100× GTP (New England Biolabs), water molecular biology grade to reach a final volume of 30 µL and incubated for 2h at 37°C then inactivated at 65°C for 20 minutes. 15 µL of this digestion product was used for sWGA with 0.1 mg/mL BSA (New England Biolabs), 1 mM dNTPs (New England Biolabs), 2.5 µM of each primer (1), 1× Phi29 reaction Buffer (New England Biolabs), 30 units of Phi29 Polymerase (New England Biolabs) and water molecular biology grade to reach a final volume of 50 µL. Amplified products were cleaned using Agencourt Ampure XP beads (Beckman Coulter).

Whole-genome sequencing

After sWGA, 250 ng of amplified DNA were used to prepare sequencing libraries using the KAPA HyperPrep Library Preparation Kit (Kapa Biosystems, Woburn, MA) following manufacturer's instructions. Covaris S220 through microTube-50 AFA Fiber Screw-Cap (Covaris®) was used to performed DNA shearing. DNA libraries were then checked for quality and quantity using Qubit® and BioAnalyzer 2100 Agilent for fragment size. Libraries were sequenced on an Illumina NextSeq 500 System using 150 bp paired-end sequencing chemistry at the GENOM'IC platform from Cochin Institute (Paris, France). Sequence data obtained from each sample was subjected to standard Illumina QC procedures. Raw reads were aligned to the PowCR01 reference genome using the BWA-mem (Burrows-Wheeler Aligner) algorithm (default parameters) (2).

Microsatellite development and analysis

All PCR products were run on 2% agarose gel electrophoresis. They were diluted before fragment analysis, if necessary, in molecular biology grade water two to ten times according to the intensity on agarose gel and run with either GeneScanTM 350 ROXTM or GeneScanTM 500 ROXTM (ThermoFisher) on an ABI 3130 genetic analyzer (ThermoFisher Scientific). Fragment analysis profiles were interpreted with the Microsatellite Analysis software on the Thermo Fisher ConnectTM website. When multiple peaks were detected, minor peaks were scored only if their height was at least one-third of the one of the dominant peak. Amplification of *P. falciparum*, *P. vivax*, *P. malariae* and *P. ovale curtisi* was tested for each marker and compared to *P. ovale wallikeri*.
